## Supplementary Figures for "CRISPR screens reveal convergent targeting strategies against evolutionarily distinct chemoresistance in cancer"

**Supplementary Information associated with this article include:**

Supplementary Figs. 1-18
Supplementary Tables 1-6 (attached datasets)

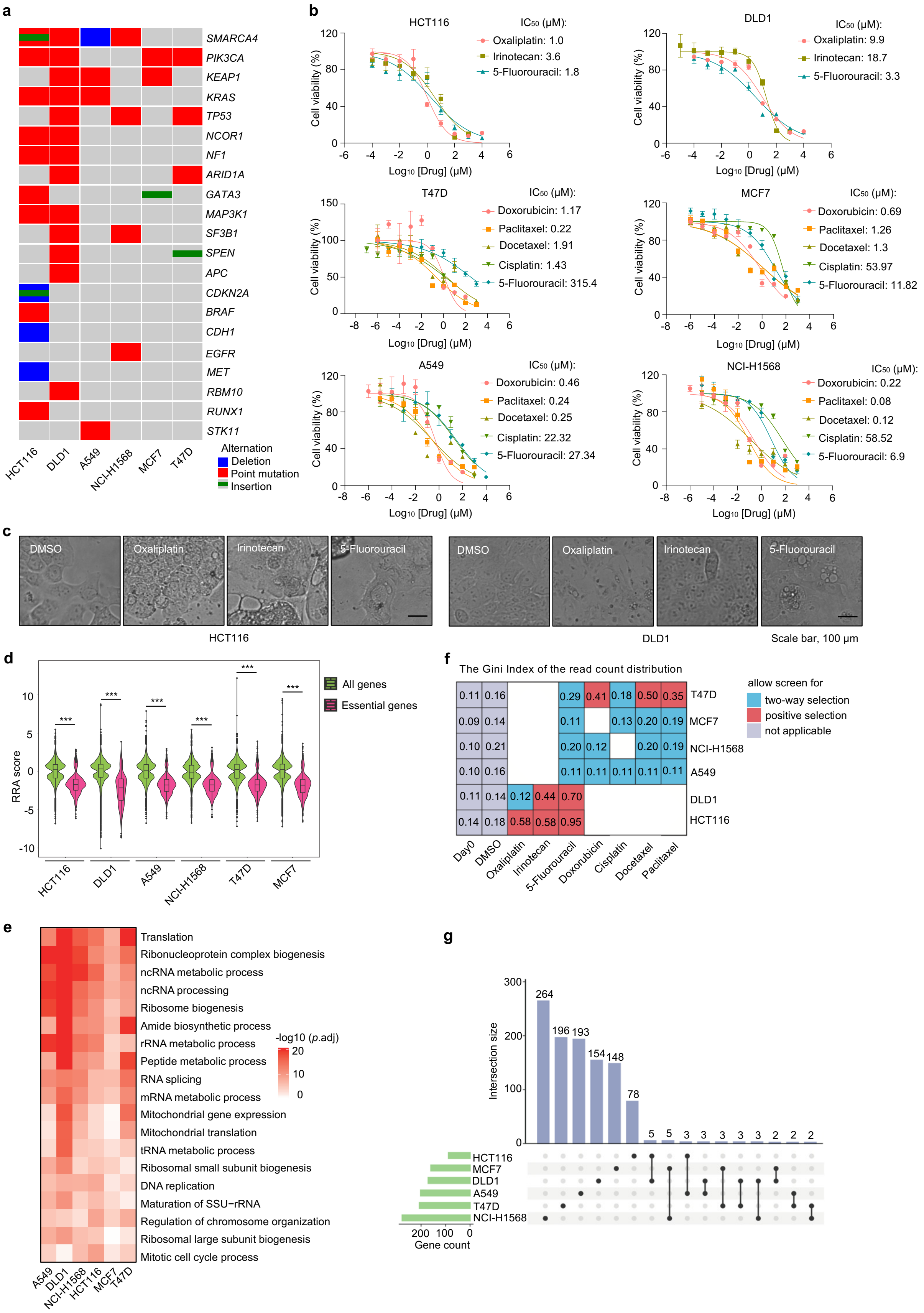

a
SMARCA4
PIK3CA
KEAP1
KRAS
TP53
NCOR1
NF1
ARID1A
GATA3
MAP3K1
SF3B1
SPEN
APC
CDKN2A
BRAF
CDH1
EGFR
MET
RBM10
RUNX1
STK11
HCT116
DLD1
A549
NCI-H1568
MCF7
T47D
Alternation
Deletion
Point mutation
Insertion
b
HCT116
IC50 (μM):
Oxaliplatin: 1.0
Irinotecan: 3.6
5-Fluorouracil: 1.8
DLD1
IC50 (μM):
Oxaliplatin: 9.9
Irinotecan: 18.7
5-Fluorouracil: 3.3
T47D
IC50 (μM):
Doxorubicin: 1.17
Paclitaxel: 0.22
Docetaxel: 1.91
Cisplatin: 1.43
5-Fluorouracil: 315.4
MCF7
IC50 (μM):
Doxorubicin: 0.69
Paclitaxel: 1.26
Docetaxel: 1.3
Cisplatin: 53.97
5-Fluorouracil: 11.82
A549
IC50 (μM):
Doxorubicin: 0.46
Paclitaxel: 0.24
Docetaxel: 0.25
Cisplatin: 22.32
5-Fluorouracil: 27.34
NCI-H1568
IC50 (μM):
Doxorubicin: 0.22
Paclitaxel: 0.08
Docetaxel: 0.12
Cisplatin: 58.52
5-Fluorouracil: 6.9
Cell viability (%)
Log10 [Drug] (μM)
c
DMSO
Oxaliplatin
Irinotecan
5-Fluorouracil
HCT116
DLD1
Scale bar, 100 μm
d
RRA score
***
All genes
Essential genes
HCT116
DLD1
A549
NCI-H1568
T47D
MCF7
f
The Gini Index of the read count distribution
0.11 0.16 0.29 0.41 0.18 0.50 0.35 T47D
0.09 0.14 0.11 0.13 0.20 0.19 MCF7
0.10 0.21 0.20 0.12 0.20 0.19 NCI-H1568
0.10 0.16 0.11 0.11 0.11 0.11 0.11 A549
0.11 0.14 0.12 0.44 0.70 DLD1
0.14 0.18 0.58 0.58 0.95 HCT116
Day0
DMSO
Oxaliplatin
Irinotecan
5-Fluorouracil
Doxorubicin
Cisplatin
Docetaxel
Paclitaxel
allow screen for
two-way selection
positive selection
not applicable
e
Translation
Ribonucleoprotein complex biogenesis
ncRNA metabolic process
ncRNA processing
Ribosome biogenesis
Amide biosynthetic process
rRNA metabolic process
Peptide metabolic process
RNA splicing
mRNA metabolic process
Mitochondrial gene expression
Mitochondrial translation
tRNA metabolic process
Ribosomal small subunit biogenesis
DNA replication
Maturation of SSU−rRNA
Regulation of chromosome organization
Ribosomal large subunit biogenesis
Mitotic cell cycle process
-log10 (p.adj)
20
10
0
A549
DLD1
NCI-H1568
HCT116
MCF7
T47D
g
Intersection size
264
196
193
154
148
78
5
5
3
3
3
3
3
2
2
2
HCT116
MCF7
DLD1
A549
T47D
NCI-H1568
Gene count

**Supplementary Fig. 1 Chemogenomic CRISPR screens.**
**a**, Mutational profile of typical cancer-related genes for indicated cancer cell lines.
**b**, Cell viability assays to determine $IC_{50}$ values of indicated drugs for corresponding cell lines. Mean ± SD with n = 3.
**c**, Cell morphology of HCT116 and DLD1 cells after chemogenomic screens before cell sample collection. Scale bar, 100 μm.
**d**, Violin plot showing the RRA score for all the genes or essential genes in the screens (DMSO condition for each cell line). Wilcoxon test, ***$p < 0.001$.
**e**, House-keeping functional terms are enriched among top 500 negatively selected genes in each screen (DMSO condition for each cell line).
**f**, The Gini Index showing the distribution of sgRNA read counts for each screen sample. A smaller value indicates more evenness of the count distribution. Significant distortion of sgRNA counts (Gini Index > 0.3) in the screen makes it inappropriate to analyze negatively selected genes.
**g**, Overlap of chemoresistance genes for the six cell lines.

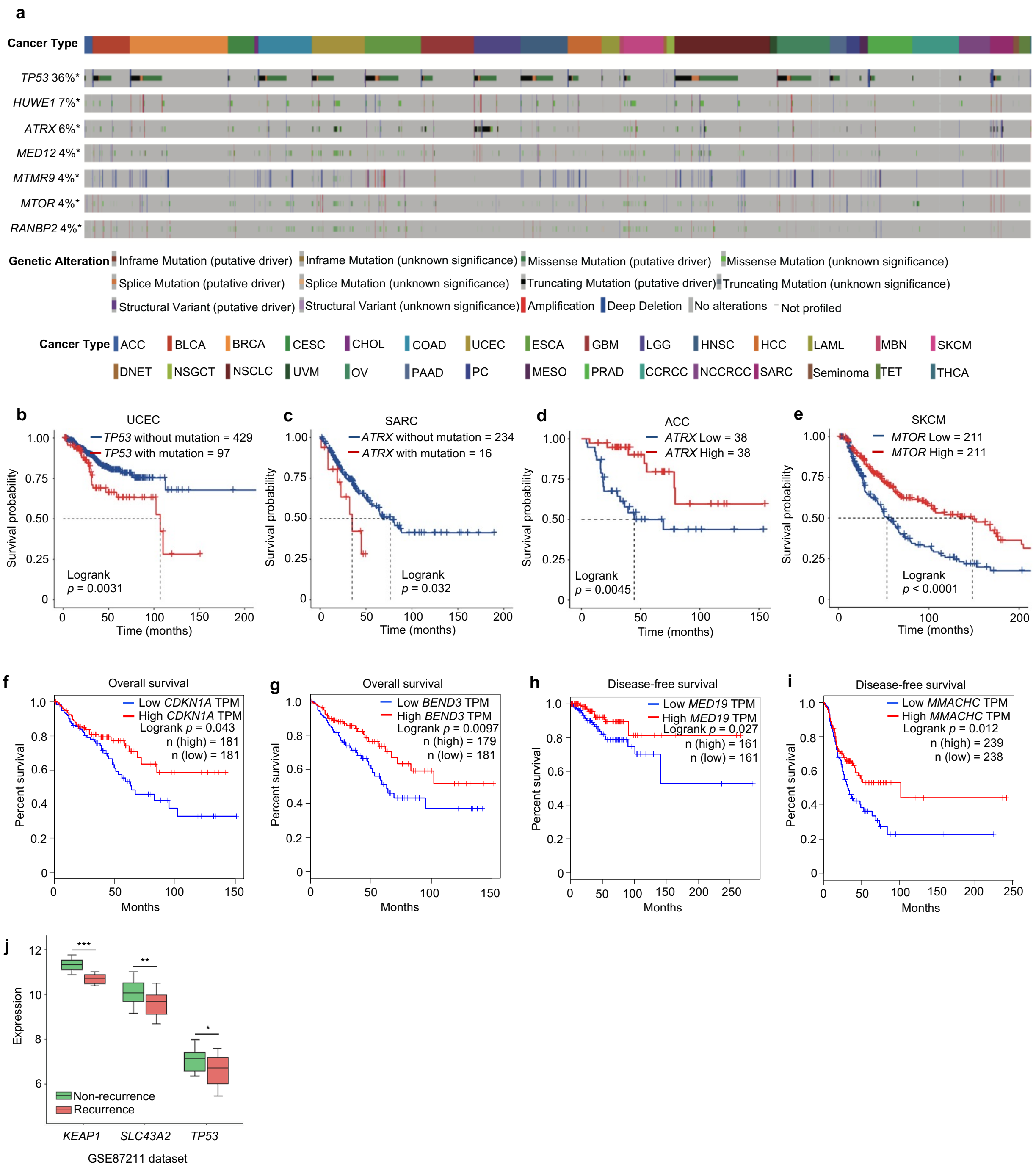

**Supplementary Fig. 2 | Identification and analysis of chemosensitizer genes.**
**a**, Overview of genetic alterations of representative chemoresistance genes in TCGA.
**b,** Overall survival analysis of Uterine Corpus Endometrial Carcinoma (UCEC) patients with *TP53* mutational status.
**c**, Overall survival analysis of Sarcoma (SARC) patients with *ATRX* mutational status.
**d**, Overall survival analysis of Adrenocortical carcinoma (ACC) patients with *ATRX* expression status.

**e**, Overall survival analysis of Skin Cutaneous Melanoma (SKCM) patients with *MTOR* expression status.

**f-g**, Overall survival analysis of colorectal cancer patients using COADREAD data from TCGA with *CDKN1A* (**f**) or *BEND3* (**g**) expression levels.

**h**, Disease-free survival analysis of breast cancer patients using BRCA data from TCGA with *MED19* expression levels.

**i**, Disease-free survival analysis of lung cancer patients using LUAD data from TCGA with *MMACHC* expression levels.

**j**, Expression levels of *KEAP1*, *TP53*, and *SLC43A2* in recurrent or non-recurrent tumors from RNA-seq data of a colorectal cancer patient cohort treated with oxaliplatin. 92 out of 203 patients in this cohort received oxaliplatin treatment and 20 patients showed recurrence after the treatment.

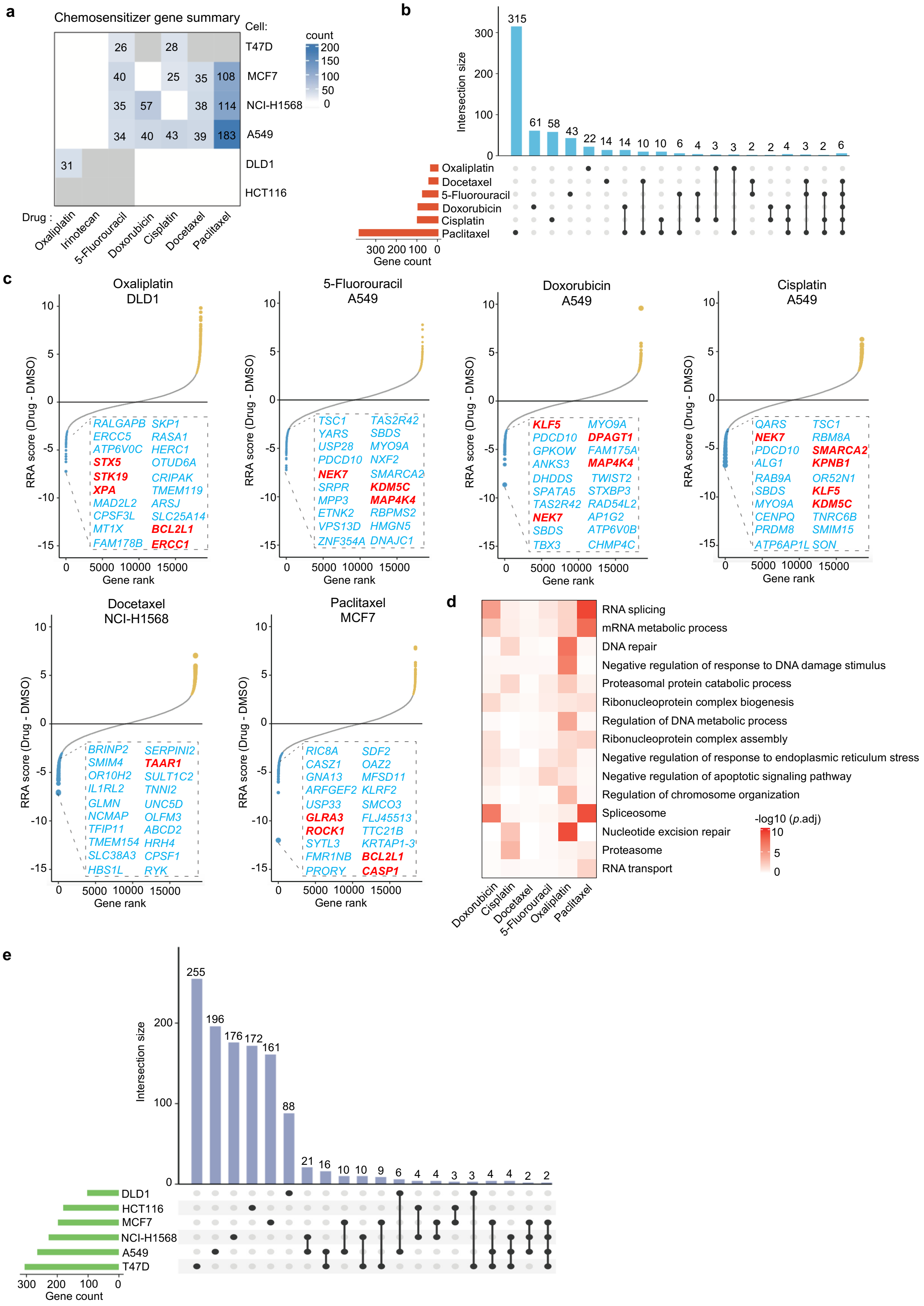
a
Chemosensitizer gene summary
Cell:
T47D
MCF7
NCI-H1568
A549
DLD1
HCT116
count
200
150
100
50
0
26
28
40
25
35
108
35
57
38
114
34
40
43
39
183
31
Drug :
Oxaliplatin
Irinotecan
5-Fluorouracil
Doxorubicin
Cisplatin
Docetaxel
Paclitaxel
b
Intersection size
315
61
58
43
22
14
14
10
10
6
4
3
3
2
2
4
3
2
6
Oxaliplatin
Docetaxel
5-Fluorouracil
Doxorubicin
Cisplatin
Paclitaxel
300 200 100 0
Gene count
c
Oxaliplatin DLD1
RRA score (Drug - DMSO)
Gene rank
RALGAPB SKP1
ERCC5 RASA1
ATP6V0C HERC1
STX5 OTUD6A
STK19 CRIPAK
XPA TMEM119
MAD2L2 ARSJ
CPSF3L SLC25A14
MT1X BCL2L1
FAM178B ERCC1
5-Fluorouracil A549
TSC1 TAS2R42
YARS SBDS
USP28 MYO9A
PDCD10 NXF2
NEK7 SMARCA2
SRPR KDM5C
MPP3 MAP4K4
ETNK2 RBPMS2
VPS13D HMGN5
ZNF354A DNAJC1
Doxorubicin A549
KLF5 MYO9A
PDCD10 DPAGT1
GPKOW FAM175A
ANKS3 MAP4K4
DHDDS TWIST2
SPATA5 STXBP3
TAS2R42 RAD54L2
NEK7 AP1G2
SBDS ATP6V0B
TBX3 CHMP4C
Cisplatin A549
QARS TSC1
NEK7 RBM8A
PDCD10 SMARCA2
ALG1 KPNB1
RAB9A OR52N1
SBDS KLF5
MYO9A KDM5C
CENPQ TNRC6B
PRDM8 SMIM15
ATP6AP1L SON
Docetaxel NCI-H1568
BRINP2 SERPINI2
SMIM4 TAAR1
OR10H2 SULT1C2
IL1RL2 TNNI2
GLMN UNC5D
NCMAP OLFM3
TFIP11 ABCD2
TMEM154 HRH4
SLC38A3 CPSF1
HBS1L RYK
Paclitaxel MCF7
RIC8A SDF2
CASZ1 OAZ2
GNA13 MFSD11
ARFGEF2 KLRF2
USP33 SMCO3
GLRA3 FLJ45513
ROCK1 TTC21B
SYTL3 KRTAP1-3
FMR1NB BCL2L1
PRORY CASP1
d
RNA splicing
mRNA metabolic process
DNA repair
Negative regulation of response to DNA damage stimulus
Proteasomal protein catabolic process
Ribonucleoprotein complex biogenesis
Regulation of DNA metabolic process
Ribonucleoprotein complex assembly
Negative regulation of response to endoplasmic reticulum stress
Negative regulation of apoptotic signaling pathway
Regulation of chromosome organization
Spliceosome
Nucleotide excision repair
Proteasome
RNA transport
-log10 (p.adj)
10
5
0
Doxorubicin
Cisplatin
Docetaxel
5-Fluorouracil
Oxaliplatin
Paclitaxel
e
Intersection size
255
196
176
172
161
88
21
16
10
10
9
6
4
4
3
3
4
4
2
2
DLD1
HCT116
MCF7
NCI-H1568
A549
T47D
300 200 100 0
Gene count

**Supplementary Fig. 3 | Identification and analysis of chemosensitizer genes.**
**a**, The number of chemosensitizer genes identified from each indicated screen.
**b**, Overlap of chemosensitizer genes for each chemotherapy drug.
**c**, Top chemosensitizer genes from representative screens for each chemotherapy drug. Genes with available targeted inhibitors are highlighted in red bold font.
**d**, Enriched functional terms from KEGG and GO for chemosensitizer genes of indicated chemotherapy drugs.
**e**, Overlap of chemosensitizer genes for the six cell lines.

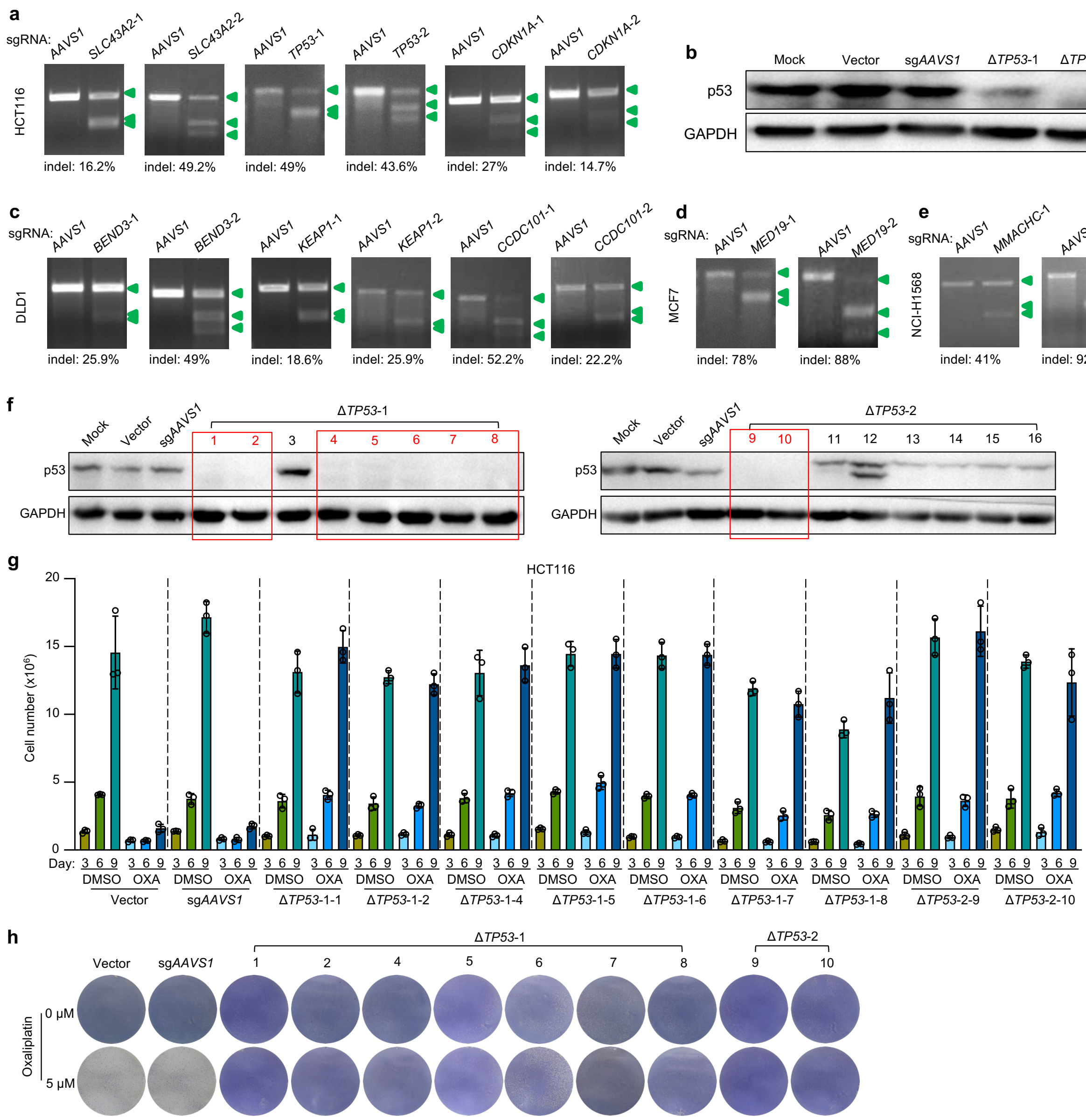


**Supplementary Fig. 4 | CRISPR-mediated knockout of individual chemoresistance genes.**

**a**, Indel assay based on T7 endonuclease I showing the effectiveness of indicated sgRNAs to generate indels at target gene loci in HCT116 cells.

**b**, Immunoblot of p53 for control and *TP53* knockout HCT116 cells.

**c**, Indel assay showing the effectiveness of indicated sgRNAs at target gene loci in DLD1 cells.

**d**, Indel assay based on T7 endonuclease I showing the effectiveness of indicated sgRNAs to generate indels at target gene loci in MCF7 cells.

**e**, Indel assay showing the effectiveness of indicated sgRNAs at target gene loci in NCI-H1568 cells.

**f**, Immunoblot of p53 for different single clones of *TP53* knockout cells. The clones with complete p53 knockout are marked in red.

**g**, Cell number quantification using a hemocytometer for indicated cells treated with DMSO or oxaliplatin. OXA, oxaliplatin. Mean ± SD with n = 3.
**h**, Crystal violet staining of indicated cells treated with oxaliplatin.

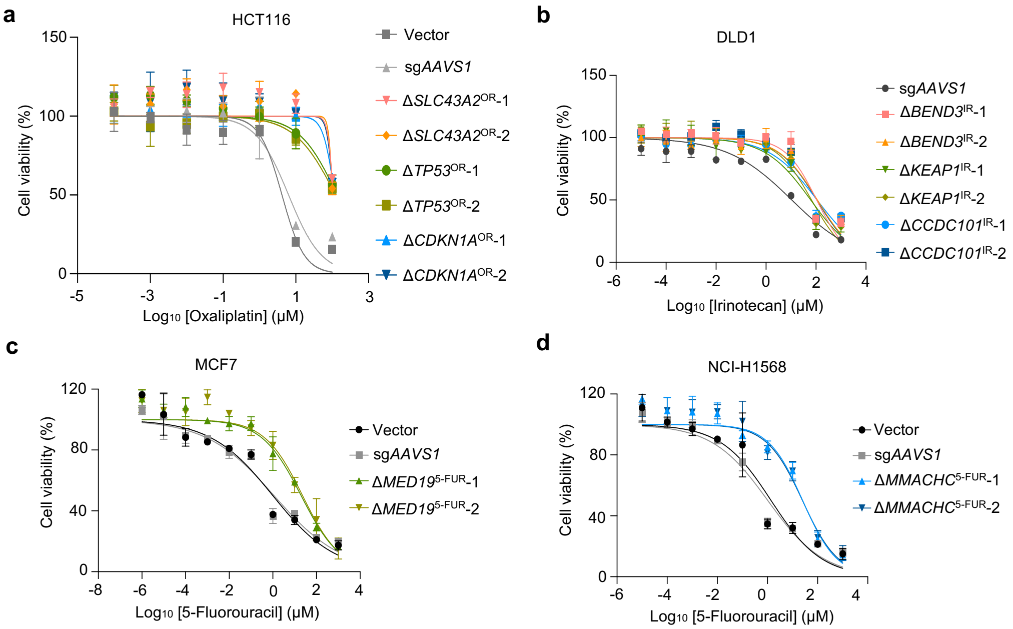


**Supplementary Fig. 5 | Cell viability assay for determining $IC_{50}$ of the drugs.**

**a**, Cell viability assay (MTT method) of different sensitive (Vector-without sgRNA, sg*AAVS1*-expressing sgRNA targeting *AAVS1* locus) and resistant HCT116 cell lines (established by indicated gene knockout and oxaliplatin selection) in response to oxaliplatin. OR, oxaliplatin resistance. -1 and -2 indicate different sgRNAs for individual gene knockout. Mean ± SD with n = 3.

**b**, Cell viability assays evaluating the response of different irinotecan-resistant DLD1 cell lines (established by individual gene knockout and irinotecan selection) to irinotecan. IR, irinotecan resistance. Mean ± SD with n = 3.

**c**, Cell viability assay of different sensitive and resistant MCF7 cell lines (established by *MED19* knockout and 5-Fluorouracil selection) in response to 5-Fluorouracil. 5-FUR, 5-Fluorouracil resistance. Mean ± SD with n = 3.

**d**, Cell viability assay of different sensitive and resistant NCI-H1568 cell lines (established by *MMACHC* knockout and 5-Fluorouracil selection) in response to 5-Fluorouracil. Mean ± SD with n = 3.

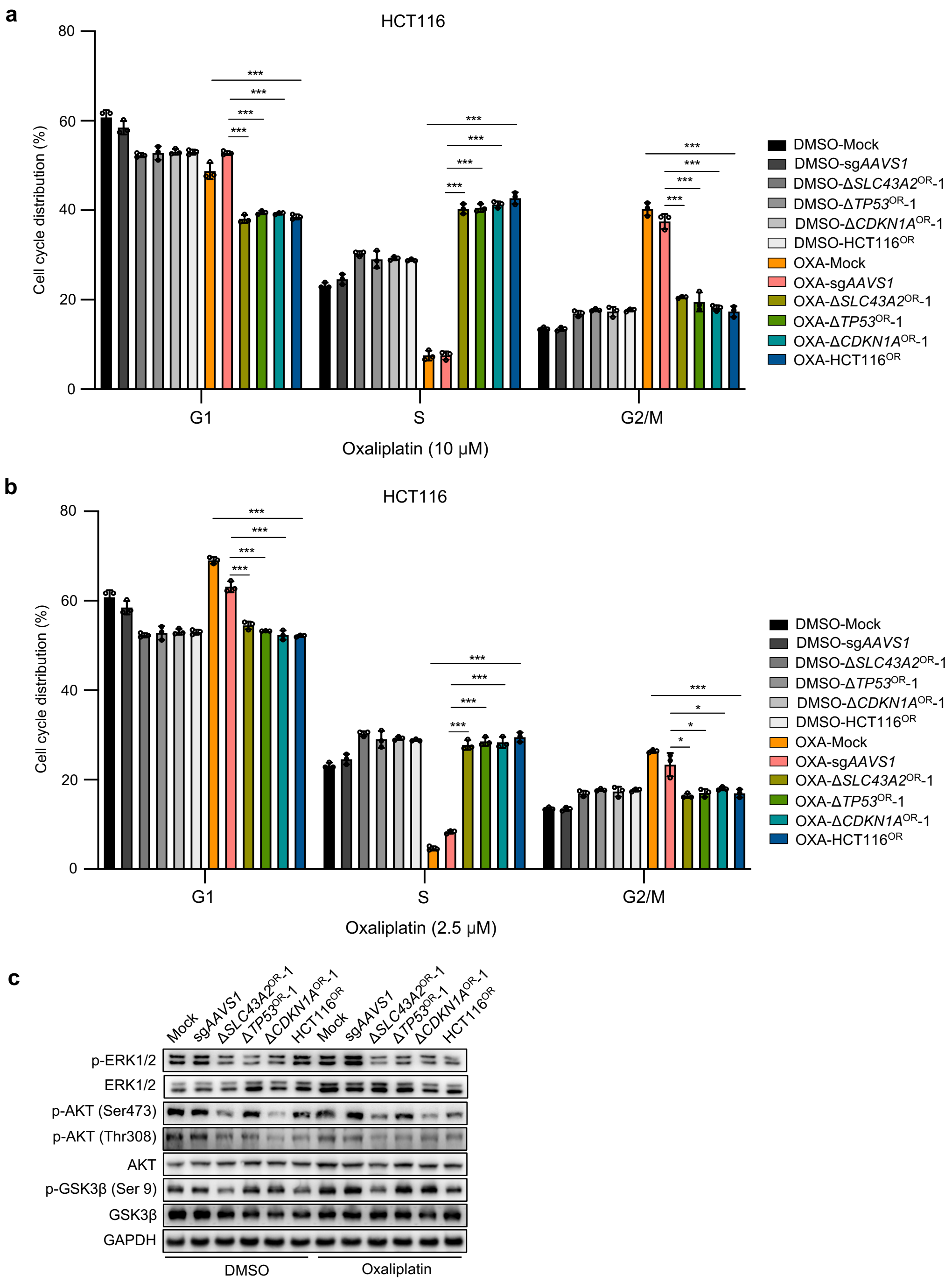


**Supplementary Fig. 6 | Additional characterization of oxaliplatin-resistant cells.**
**a-b**, Cell cycle analysis by propidium iodide (PI) staining of indicated cell lines in the absence or presence of 10 µM (**a**) or 2.5 µM (**b**) oxaliplatin for 48 h. OXA, oxaliplatin. Mean ± SD with n = 3. Unpaired t test, *$p < 0.05$ and ***$p < 0.001$.
**c**, Immunoblot analysis of indicated proteins for multiple oxaliplatin-sensitive or -resistant HCT116 cell lines in the absence or presence of 5 µM oxaliplatin for 72 h.

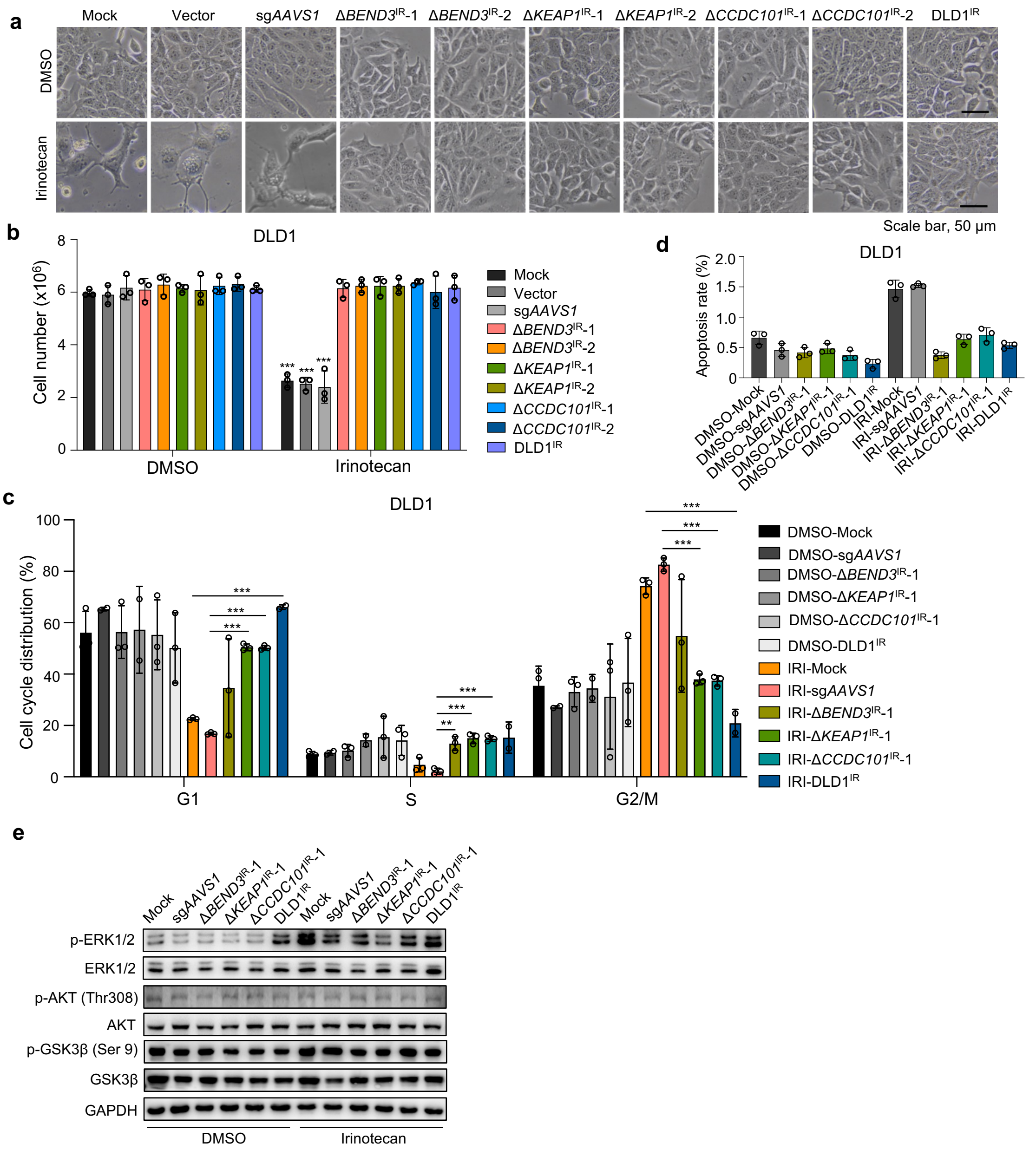


**Supplementary Fig. 7 | Cellular and molecular features of irinotecan-resistant cells.**

**a**, Cell morphology of multiple irinotecan-sensitive (Mock, Vector and sg*AAVS1* groups) and -resistant cell lines in the absence or presence of irinotecan. Scale bar, 50 µm.

**b**, Cell number quantification of indicated irinotecan-sensitive and -resistant cell lines after 7 days of irinotecan treatment. Mean ± SD with n = 3. Unpaired t test (irinotecan vs. DMSO), $***p < 0.001$.

**c**, Cell cycle analysis by PI staining of indicated cell lines in the absence or presence of 10 µM irinotecan for 24 h. IRI, irinotecan. Mean ± SD with n = 3. Unpaired t test, $**p < 0.01$ and $***p < 0.001$.

**d**, Apoptosis analysis by PI and Hoechst staining of indicated cell lines in the absence or presence of 10 µM irinotecan for four days. Mean ± SD with n = 3.

**e**, Immunoblot analysis of indicated proteins for multiple irinotecan-sensitive or -resistant DLD1 cell lines in the absence or presence of 10 μM irinotecan for 72 h.

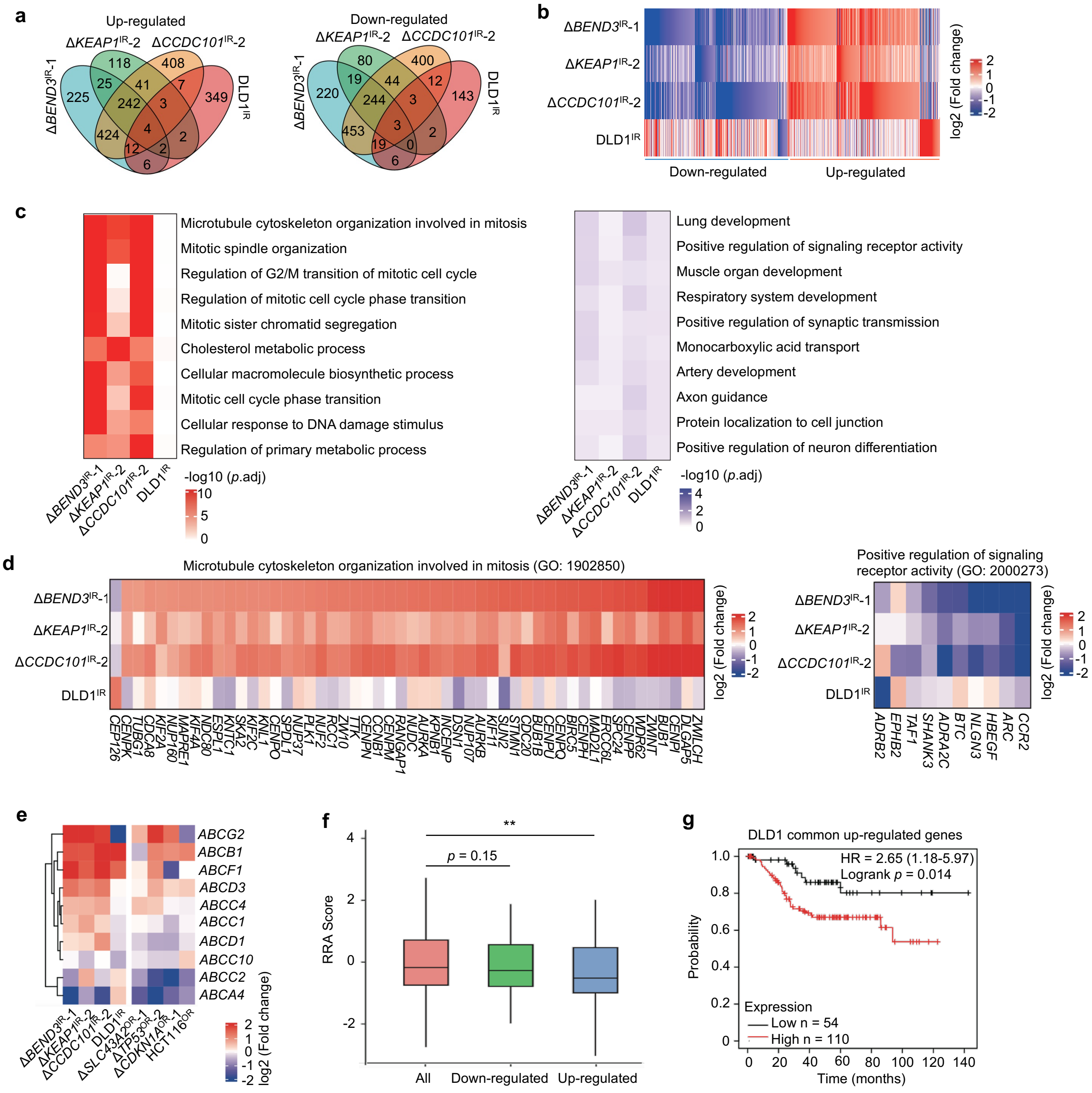

**Supplementary Fig. 8 | Gene expression profiles of irinotecan-resistant cells.**

**a**, Venn diagram of differentially expressed genes (Δ*BEND3*$^{IR}$-1, Δ*KEAP1*$^{IR}$-2 or Δ*CCDC101*$^{IR}$-2 vs. sg*AAVS1*; DLD1$^{IR}$ vs. untreated Mock) in four indicated irinotecan-resistant DLD1 cell lines determined by RNA-seq analysis.

**b**, Heatmap showing differentially expressed genes in four indicated irinotecan-resistant DLD1 cell lines.

**c**, Functional enrichment analysis showing the top enriched terms among up-regulated (left) or down-regulated (right) genes in four DLD1 irinotecan-resistant cell lines relative to -sensitive control cells.

**d**, Highlight of individual genes within indicated functional terms across the four irinotecan-resistant cell lines.

**e**, Expression change of typical ABC transporter genes associated with multidrug resistance.

**f**, RRA Score for commonly up- or down-regulated genes in Δ*BEND3*$^{IR}$, Δ*KEAP1*$^{IR}$ and Δ*CCDC101*$^{IR}$ resistance models during DLD1-irinotecan chemoresistance screen.
**g**, Survival analysis of commonly up-regulated genes in Δ*BEND3*$^{IR}$, Δ*KEAP1*$^{IR}$ and Δ*CCDC101*$^{IR}$ resistance models in a colorectal cancer patient cohort (GSE17538).

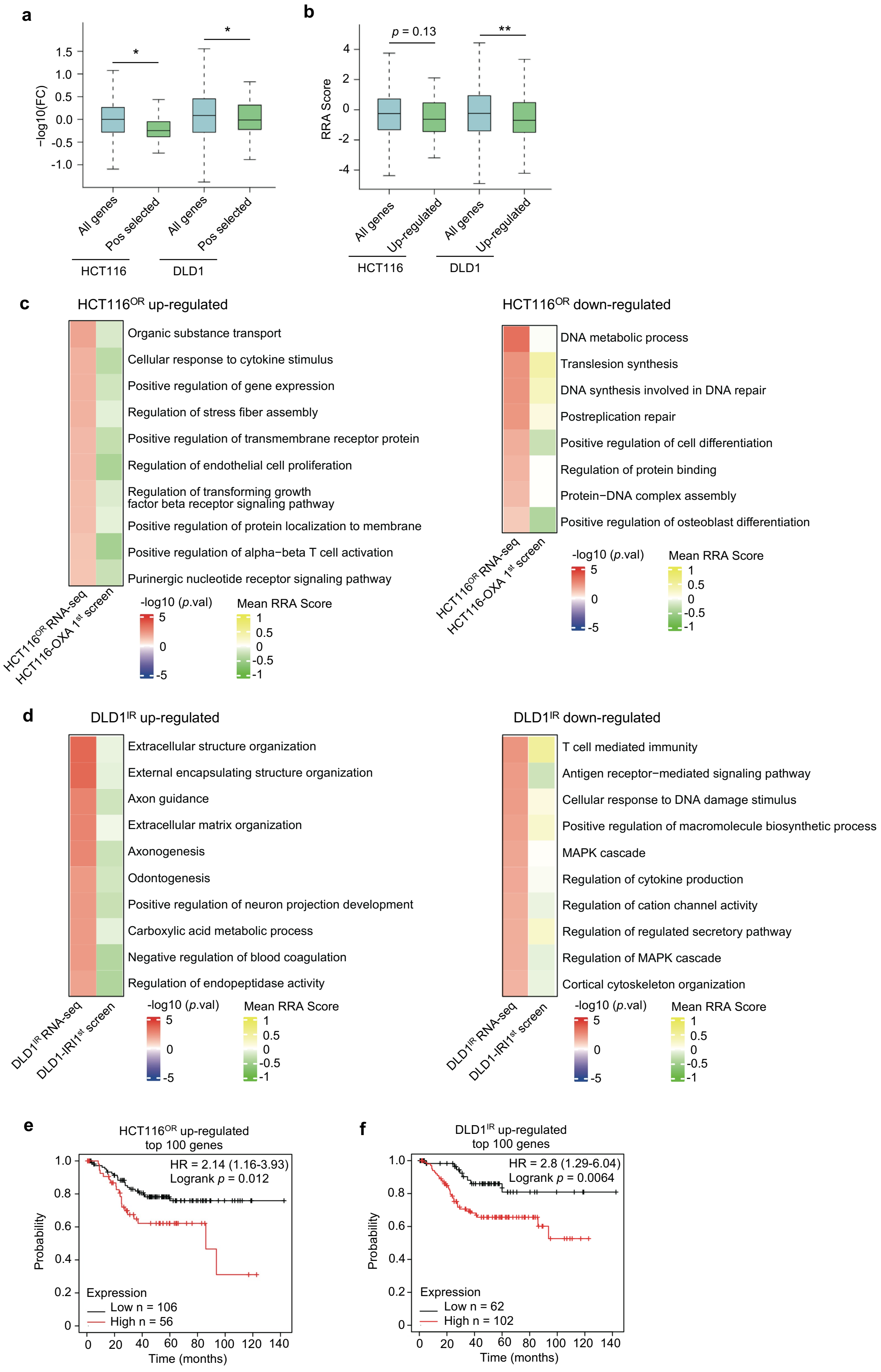
a
-log10(FC)
1.5
1.0
0.5
0.0
-0.5
-1.0
*
*
All genes
Pos selected
All genes
Pos selected
HCT116
DLD1
b
RRA Score
p = 0.13
**
4
2
0
-2
-4
All genes
Up-regulated
All genes
Up-regulated
HCT116
DLD1
c
HCT116OR up-regulated
Organic substance transport
Cellular response to cytokine stimulus
Positive regulation of gene expression
Regulation of stress fiber assembly
Positive regulation of transmembrane receptor protein
Regulation of endothelial cell proliferation
Regulation of transforming growth factor beta receptor signaling pathway
Positive regulation of protein localization to membrane
Positive regulation of alpha−beta T cell activation
Purinergic nucleotide receptor signaling pathway
HCT116OR RNA-seq
HCT116-OXA 1st screen
-log10 (p.val)
5
0
-5
Mean RRA Score
1
0.5
0
-0.5
-1
HCT116OR down-regulated
DNA metabolic process
Translesion synthesis
DNA synthesis involved in DNA repair
Postreplication repair
Positive regulation of cell differentiation
Regulation of protein binding
Protein−DNA complex assembly
Positive regulation of osteoblast differentiation
HCT116OR RNA-seq
HCT116-OXA 1st screen
d
DLD1IR up-regulated
Extracellular structure organization
External encapsulating structure organization
Axon guidance
Extracellular matrix organization
Axonogenesis
Odontogenesis
Positive regulation of neuron projection development
Carboxylic acid metabolic process
Negative regulation of blood coagulation
Regulation of endopeptidase activity
DLD1IR RNA-seq
DLD1-IRI1st screen
DLD1IR down-regulated
T cell mediated immunity
Antigen receptor−mediated signaling pathway
Cellular response to DNA damage stimulus
Positive regulation of macromolecule biosynthetic process
MAPK cascade
Regulation of cytokine production
Regulation of cation channel activity
Regulation of regulated secretory pathway
Regulation of MAPK cascade
Cortical cytoskeleton organization
DLD1IR RNA-seq
DLD1-IRI1st screen
e
HCT116OR up-regulated top 100 genes
HR = 2.14 (1.16-3.93)
Logrank p = 0.012
Probability
Expression
Low n = 106
High n = 56
Time (months)
0 20 40 60 80 100 120 140
f
DLD1IR up-regulated top 100 genes
HR = 2.8 (1.29-6.04)
Logrank p = 0.0064
Probability
Expression
Low n = 62
High n = 102
Time (months)
0 20 40 60 80 100 120 140

**Supplementary Fig. 9 | Relationship between chemoresistant genes from CRISPR screen and differential genes from RNA-seq in HCT116$^{OR}$ or DLD1$^{IR}$ resistant models.**

**a**, The expression status of positively selected gene hits during corresponding chemoresistance CRISPR screen in RNA-seq data of HCT116$^{OR}$ and DLD1$^{IR}$ resistance models.

**b**, The RRA score of significantly up-regulated genes in HCT116$^{OR}$ and DLD1$^{IR}$ resistance models during HCT116-oxaliplatin or DLD1-irinotecan chemoresistant screen.

**c**, The top enriched functional terms for up-regulated or down-regulated genes in HCT116$^{OR}$ resistance model and their mean RRA scores for the genes in corresponding terms during HCT116-oxaliplatin chemoresistance screen.

**d**, The top enriched functional terms for up-regulated or down-regulated genes in DLD1$^{IR}$ resistance model and their mean RRA scores for the genes in corresponding terms during DLD1-irinotecan chemoresistance screen.

**e-f**, Survival analysis of top 100 up-regulated genes in HCT116$^{OR}$ (**e**) or DLD1$^{IR}$ (**f**) resistance models in a colorectal cancer patient cohort (GSE17538).

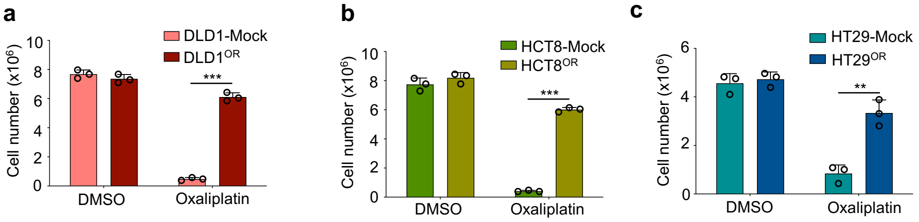


**Supplementary Fig. 10 | Establishment of additional oxaliplatin-resistant lines with various genetic backgrounds by traditional drug pulsing and clonal selection method.**
**a**-**c**, Cell growth response to oxaliplatin for indicated parental cells and resistant cells established by "randomly" clonal selection with gradual increase of oxaliplatin treatment. Cell number was quantified after treatment of DLD1 cells with 80 μM oxaliplatin for 7 days (**a**), HCT8 cells with 50 μM oxaliplatin for 7 days (**b**) and HT29 cells with 35 μM oxaliplatin for 10 days (**c**), respectively. Mean ± SD with n = 3. Unpaired t test, $**p < 0.01$ and $***p < 0.001$.

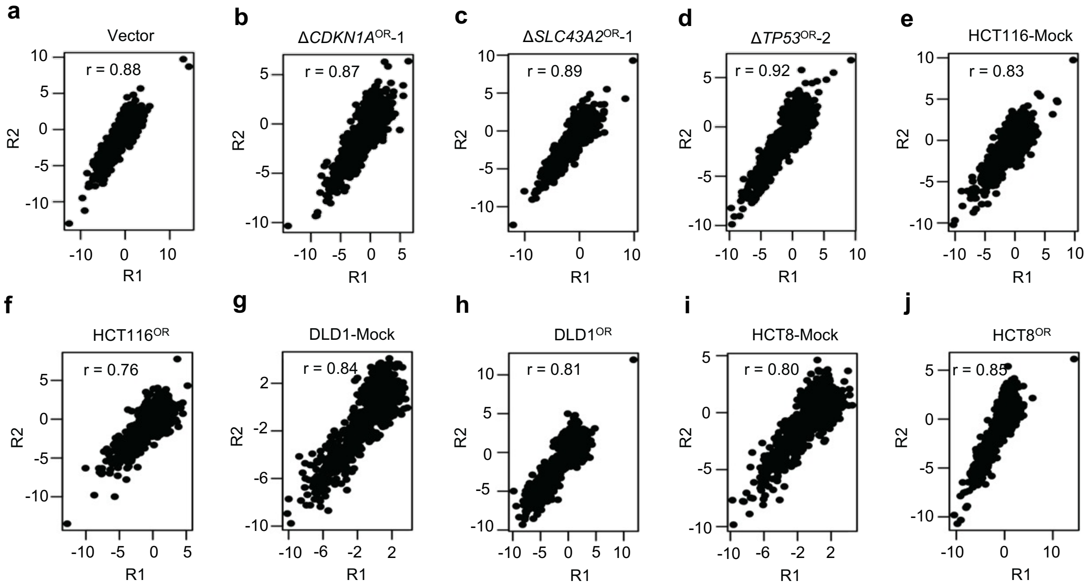


**Supplementary Fig. 11 | Replicate correlation for indicated second-round CRISPR screen samples.**

**a-j**, Correlation of RRA scores between the two biological replicates (R1 and R2) across all the indicated conditions during second-round druggable CRISPR screens against oxaliplatin resistance.

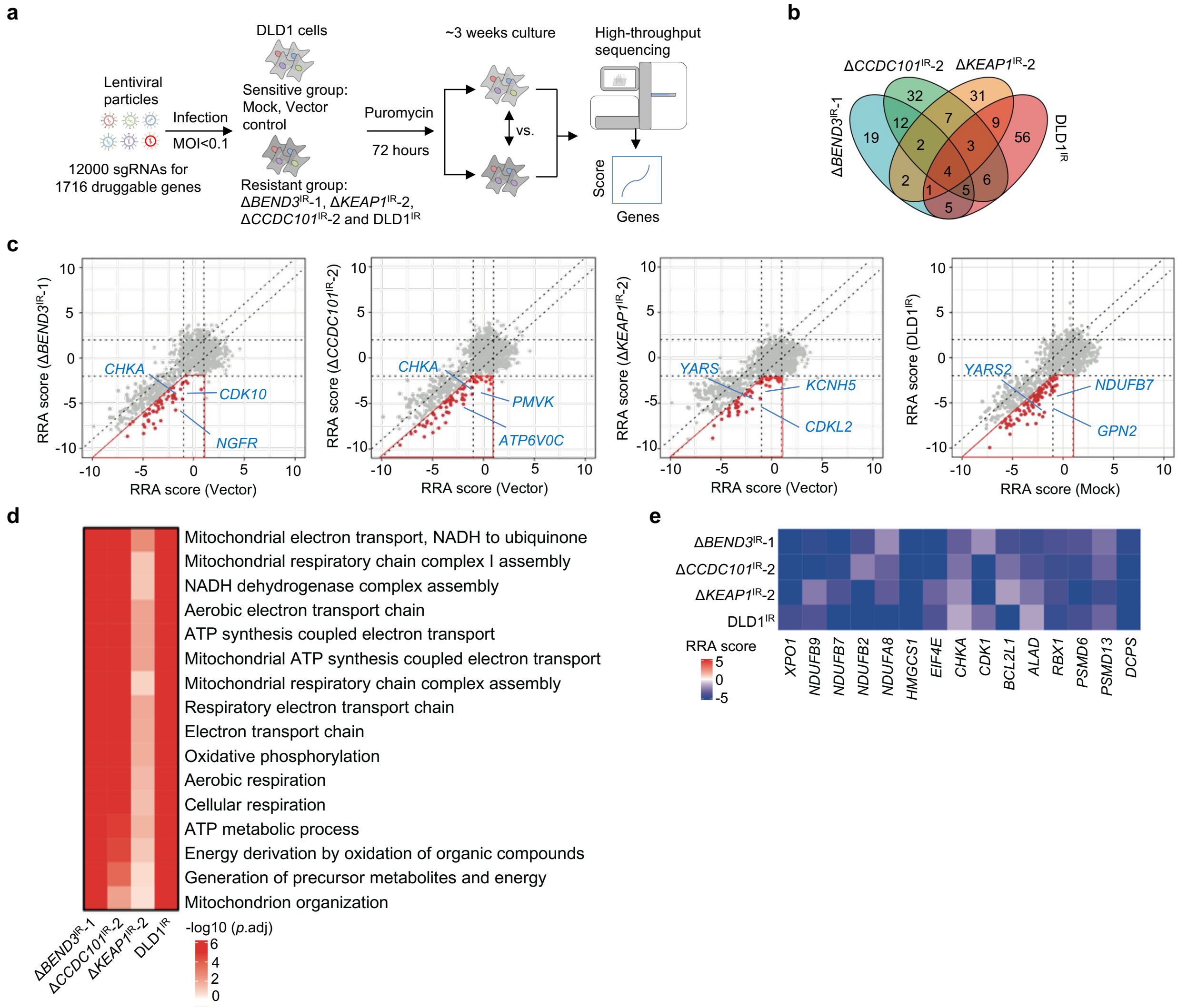


**Supplementary Fig. 12 | Interrogation of druggable targets against irinotecan resistance by second-round CRISPR screens.**

**a**, Workflow of identification of potential targets against irinotecan resistance using druggable gene CRISPR knockout screens.

**b**, Venn diagram of preferably druggable gene hits (Δ*BEND3*$^{IR}$-1, Δ*KEAP1*$^{IR}$-2 or Δ*CCDC101*$^{IR}$-2 vs. Vector; DLD1$^{IR}$ vs. untreated Mock) in four irinotecan-resistant DLD1 cell lines by second-round CRISPR screens.

**c**, The RRA scores of all the tested genes for irinotecan-sensitive control or -resistance cells in each comparison. The genes in red box are preferential targets against resistance. Several representative genes are highlighted.

**d**, The top enriched functional terms for druggable gene sets against irinotecan identified from second-round CRISPR screens in four resistant cell lines.

**e**, Heatmap showing the RRA scores for the top consensus druggable targets across all the four irinotecan-resistant cell lines.

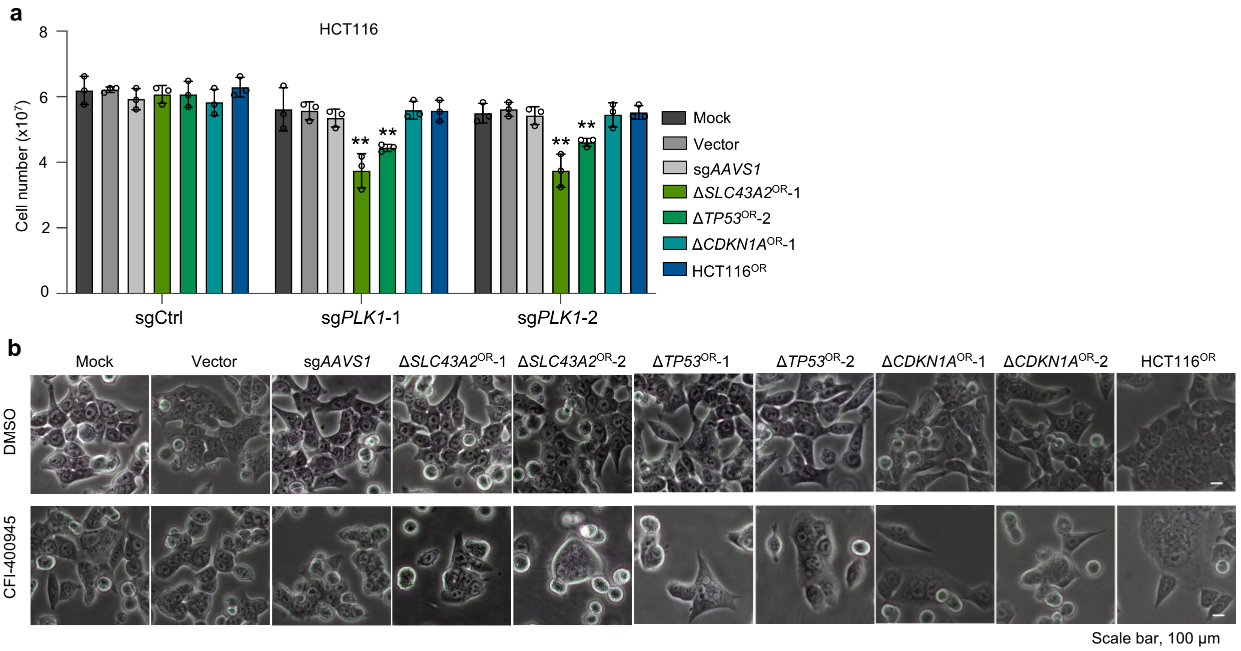


**Supplementary Fig. 13 | Cell growth effect of *PLK1* knockout and cellular morphology upon PLK4 inhibitor treatment for oxaliplatin-resistant cells.**

**a**, Cell growth effect of *PLK1* knockout by two independent sgRNAs in oxaliplatin-sensitive or -resistant cell lines. Mean ± SD with n = 3. Unpaired t test (Δ*SLC43A2*$^{OR}$, Δ*TP53*$^{OR}$ or Δ*CDKN1A*$^{OR}$ vs. sg*AAVS1*; HCT116$^{OR}$ vs. Mock), $**p < 0.01$.

**b**, Cell morphology of multiple oxaliplatin-sensitive (Mock, Vector and sg*AAVS1* groups) and -resistant cell lines in the absence or presence of 12 nM CFI-400945 for one week. Scale bar, 100 μm.

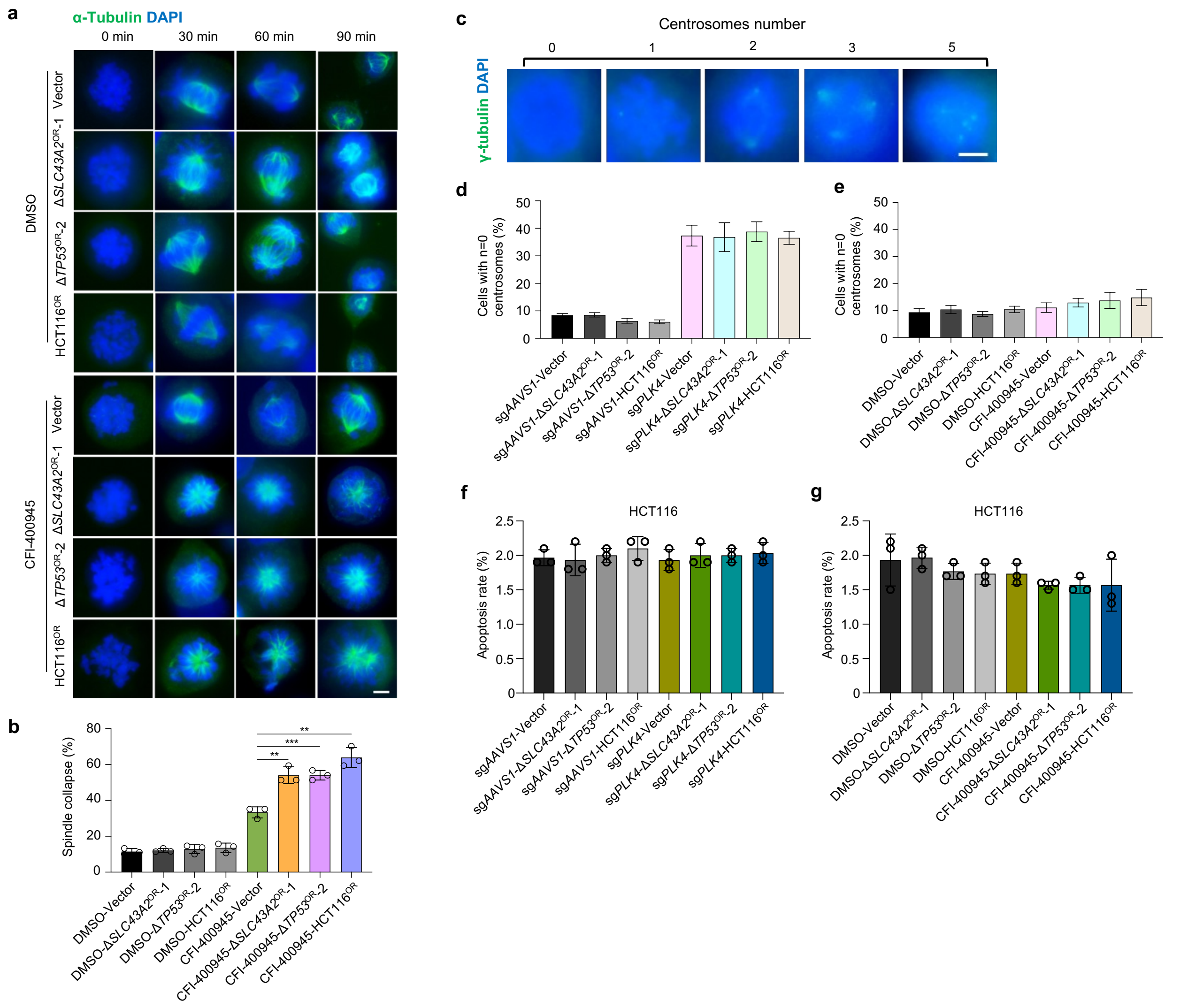


**Supplementary Fig. 14 | Targeting PLK4 in oxaliplatin-resistant cells.**

**a**, Treatment with CFI-400945 results in spindle collapse, folding and slippage in oxaliplatin-resistant cells. Scale bar, 5 µm.

**b**, Quantification of spindle collapse for (**a**). Mean ± SD with n = 3, 100-200 cells were analyzed per independent experiment. Unpaired t test, **p* < 0.05 and ***p* < 0.01

**c**, Representative images for centrosome quantification by γ-tubulin staining.

**d-e**, Distribution of cell populations with zero centrosome number before or after *PLK4* knockout (**d**) or CFI- 400945 treatment (**e**) for oxaliplatin-sensitive and -resistant cells. Mean ± SD with n = 3, 100-200 cells were analyzed per independent experiment. Unpaired t test, **p* < 0.05 and ***p* < 0.01.

**f**, Apoptosis assay by PI and Annexin V staining showing that knockout of *PLK4* does not affect apoptosis of oxaliplatin-sensitive or -resistant cell lines. Mean ± SD with n = 3.

**g**, Apoptosis assay by PI and Annexin V staining showing that CFI-400945 does not affect apoptosis of oxaliplatin-sensitive or -resistant cell lines. Mean ± SD with n = 3.

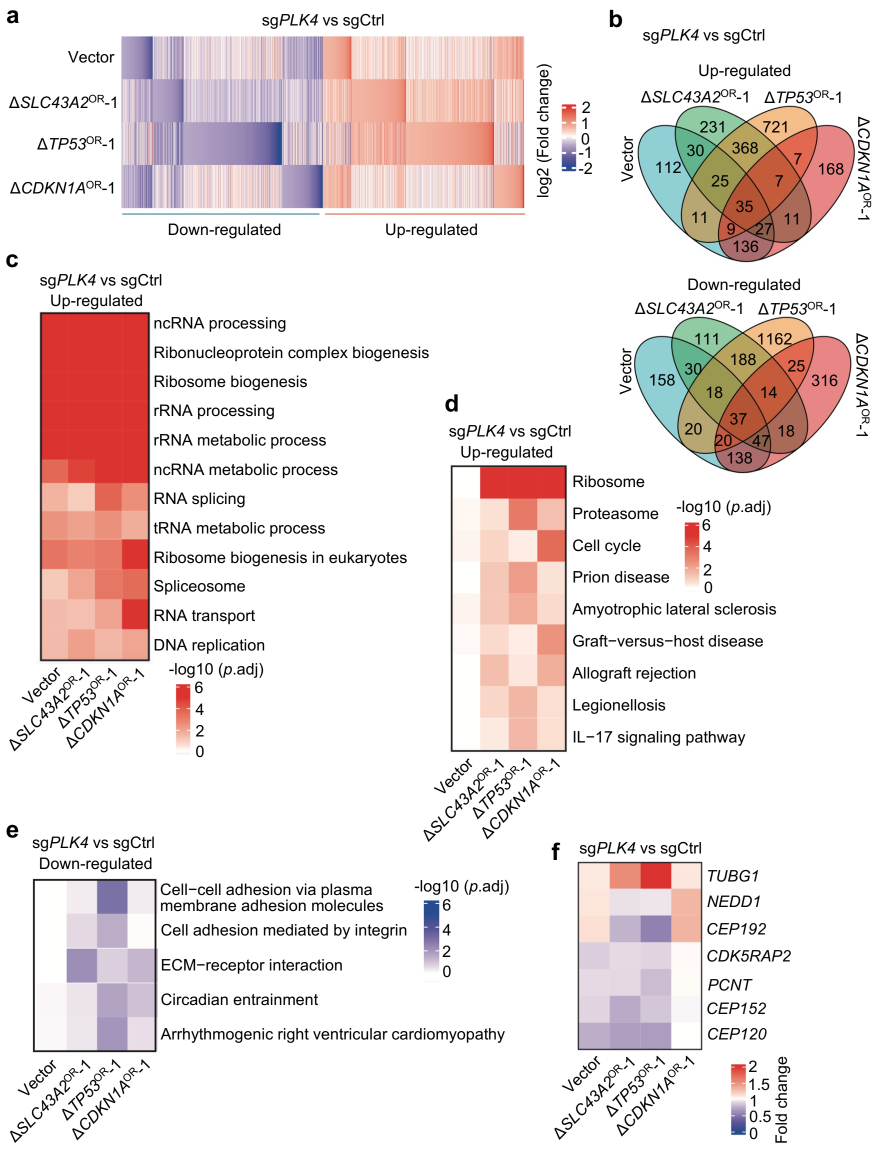


**Supplementary Fig. 15 | RNA-seq analysis of PLK4 target genes in oxaliplatin-sensitive or -resistant cells.**

**a**, Heatmap showing *PLK4* downstream target genes of indicated oxaliplatin-sensitive (Vector) and -resistant cells determined by RNA-seq.

**b**, Venn diagram of *PLK4* target genes in four indicated cell lines.

**c**, Functional terms enriched among up-regulated *PLK4* target genes for each indicated cell line.

**d**-**e**, Functional terms specifically enriched among up-regulated genes (**d**) or down-regulated genes (**e**) upon *PLK4* knockout in oxaliplatin-resistant cell lines but not in sensitive cells (Vector).

**f**, The expression change of typical pericentriolar material (PCM) scaffolding genes upon *PLK4* knockout in each indicated cell line.

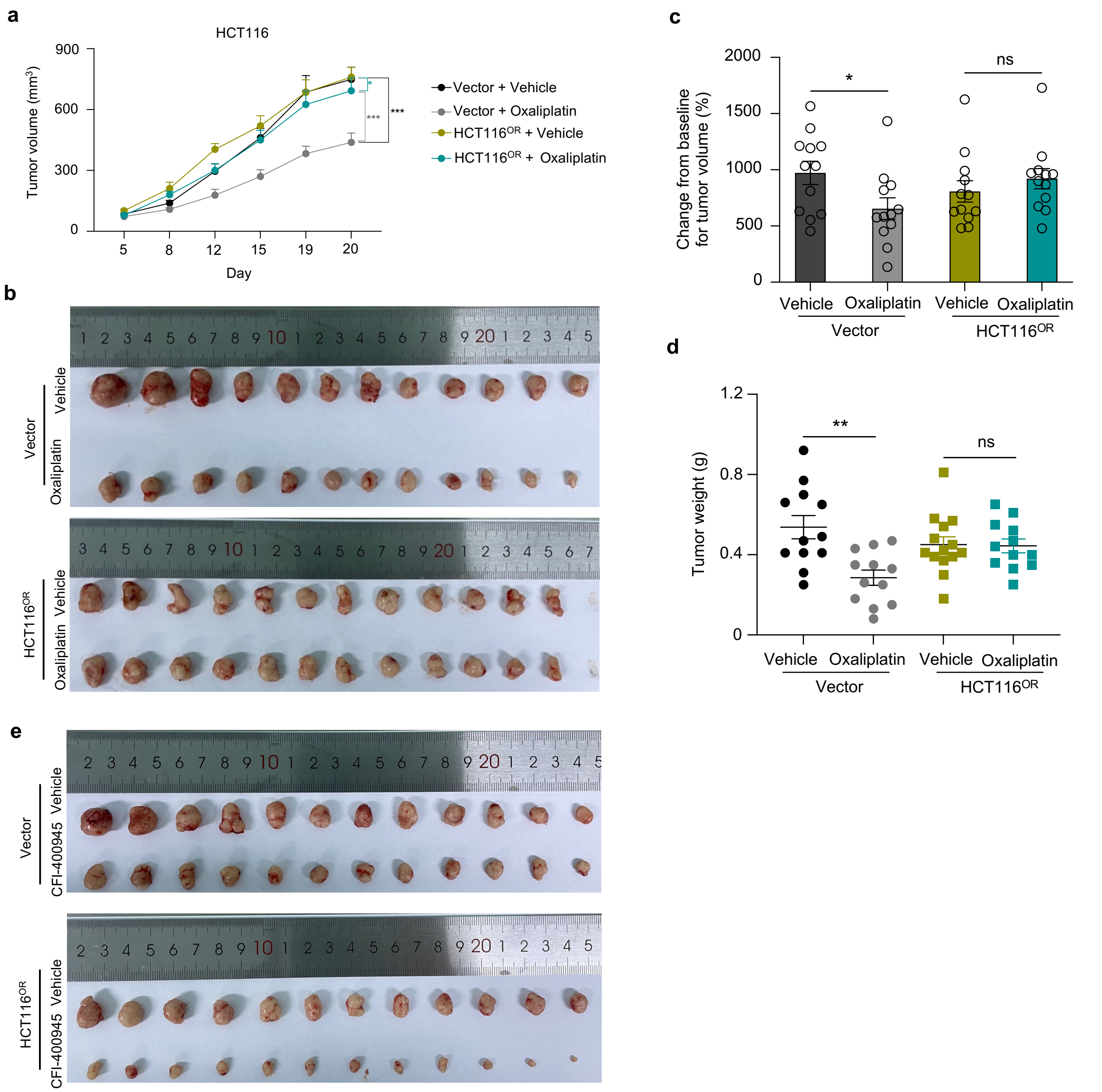


**Supplementary Fig. 16 | Validation of oxaliplatin resistance by xenograft model in mice.**
**a**, Tumor volume measured at indicated time points after xenograft implantation for oxaliplatin-sensitive (Vector) or -resistant (HCT116$^{OR}$) xenograft treated with vehicle or oxaliplatin. (n=12, number of tumor). Mean ± SEM. Two-way ANOVA, ***$p < 0.001$.
**b**, Photos of tumors collected from oxaliplatin-sensitive or -resistance xenograft bearing mice with vehicle or oxaliplatin treatment at the experimental endpoint.
**c**-**d**, Relative tumor volume (compared to day 0) (**c**) or tumor weight (**d**) of mouse xenografts for indicated HCT116 cells measured at Day 20 post implantation. (n=12, number of tumor). Mean ± SEM. Unpaired t test, *$p < 0.05$ and **$p < 0.01$. ns, non-significant.
**e**, Photos of tumors collected from oxaliplatin-sensitive or -resistance xenograft bearing mice with vehicle or CFI-400549 treatment at the experimental endpoint.

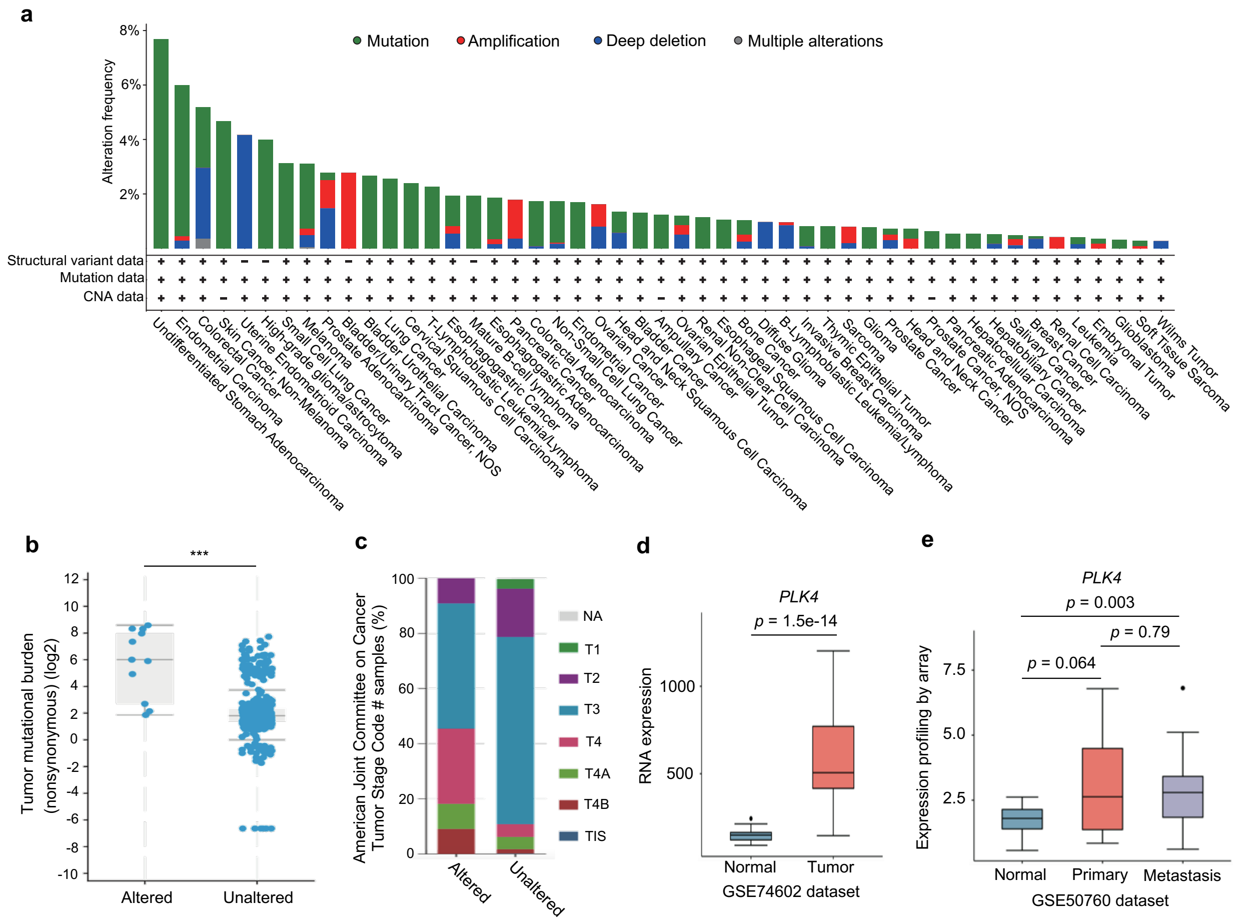


**Supplementary Fig. 17 | Clinical relevance of PLK4.**

**a**, Overview of *PLK4* alterations across multiple cancers from cBioPortal.

**b**, Colorectal cancers with mutated *PLK4* tend to have significantly higher tumor mutational burden according to TCGA data from cBioPortal.

**c**, *PLK4* mutation is positively associated with advanced progression of colorectal cancers.

**d**, RNA expression of *PLK4* in GSE74602 dataset. Significance was calculated by the Wilcoxon test.

**e**, RNA expression of *PLK4* in GSE50760 dataset. Significance was calculated by the Wilcoxon test.

**a**

| Organoid number | MLH1 | MSH2 | MSH6 | PMS2 | Ki-67 | CDX2 | HER2 | CK7 | CK20 | BRAF |
|---|---|---|---|---|---|---|---|---|---|---|
| C-1 | + | + | + | + | unknown | unknown | unknown | unknown | unknown | unknown |
| C-2 | + | + | + | + | 80%+ | + | - | - | + | - |
| C-3 | + | + | + | + | 70%+ | + | - | - | + | - |
| C-4 | + | + | + | + | 40%+ | + | - | - | + | - |
| C-5 | + | + | + | + | 80%+ | + | - | - | + | - |
| C-6 | + | + | + | + | 60%+ | - | - | - | + | + |
| C-7 | + | + | + | + | 80%+ | + | - | - | + | - |

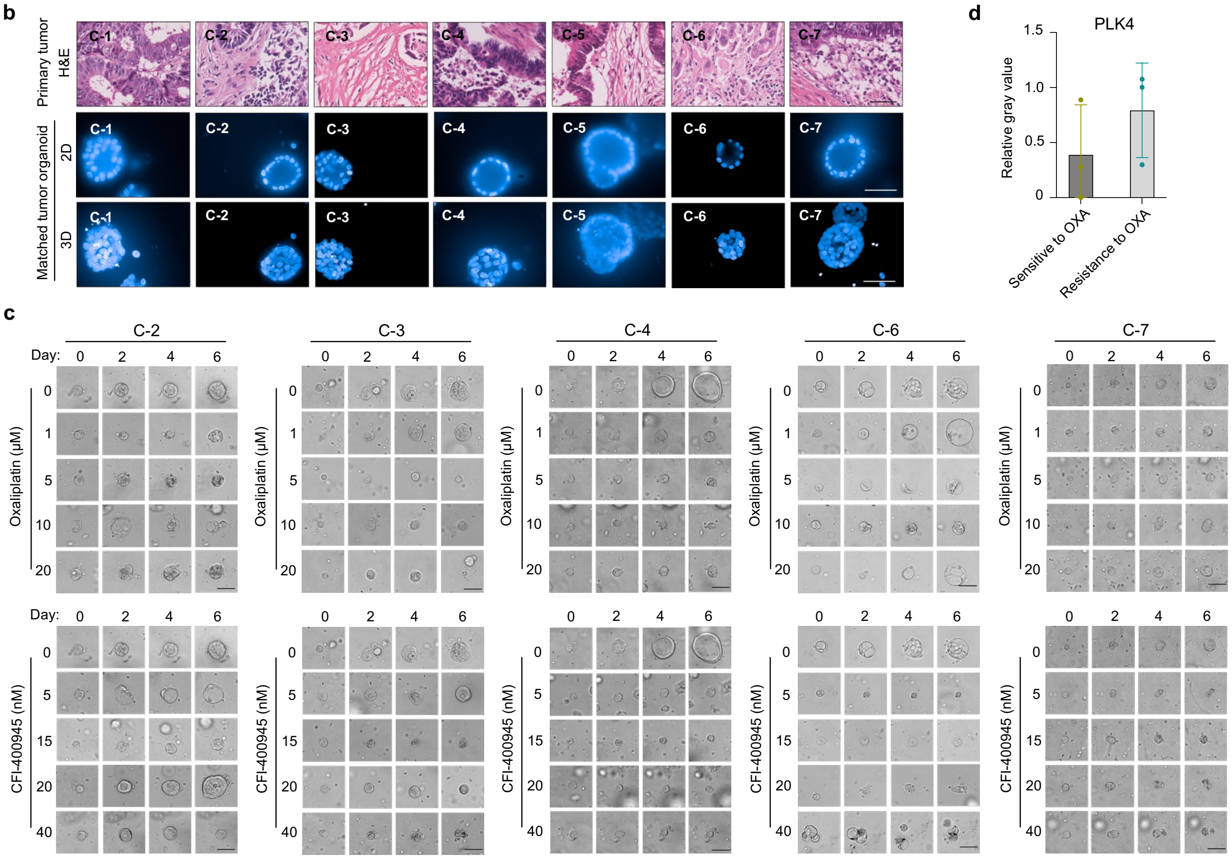


**Supplementary Fig. 18 | Characterization and drug sensitivity test of patient-derived organoids.**

**a**, Molecular characterization for several clinical biomarkers in tumors for deriving organoids.

**b**, Images showing H&E staining of primary tumors and Hoechst 33342-stained tumor-derived organoids. Scale bar, 50 μm.

**c**, Representative images showing organoid growth in response to oxaliplatin or CFI-400945 for indicated organoids. Scale bar, 50 μm.

**d**, Comparison of relative protein expression of PLK4 in oxaliplatin-resistant tumor tissues (C-5, C-6 and C-7) vs. in oxaliplatin-sensitive tumor tissues (C-1, C-2 and C-3) according to immunoblot image in Fig. 7d.
